# Propofol disrupts alpha dynamics in distinct thalamocortical networks underlying sensory and cognitive function during loss of consciousness

**DOI:** 10.1101/2022.04.05.487190

**Authors:** Veronica S. Weiner, David W. Zhou, Pegah Kahali, Emily P. Stephen, Robert A. Peterfreund, Linda S. Aglio, Michele D. Szabo, Emad N. Eskandar, Andrés F. Salazar-Gomez, Aaron L. Sampson, Sydney S. Cash, Emery N. Brown, Patrick L. Purdon

**Affiliations:** Department of Brain and Cognitive Sciences, Massachusetts Institute of Technology, Cambridge, MA, USA; Picower Institute for Learning and Memory, Massachusetts Institute of Technology, Cambridge, MA, USA; Department of Anesthesia, Critical Care and Pain Medicine, Massachusetts General Hospital, Boston, MA, USA; Center for Neurotechnology and Recovery, Department of Neurology, Massachusetts General Hospital, Boston, MA, USA; Harvard Medical School, Boston, MA, USA; Department of Anesthesiology, Perioperative and Pain Medicine, Brigham and Women’s Hospital, Boston, MA, USA; Department of Neurological Surgery, Massachusetts General Hospital, Boston, MA, USA; Division of Health Sciences and Technology, Harvard Medical School/Massachusetts Institute of Technology, Cambridge, MA, USA; Institute of Medical Engineering and Sciences, Massachusetts Institute of Technology, Cambridge, MA, USA

**Author notes:** Department of Neurology, Keck Medical Center, University of Southern California, Los Angeles, CA, USA. Department of Math and Statistics, Boston University, Boston, MA, USA. Department of Neurological Surgery, Albert Einstein College of Medicine—Montefiore Medical Center, Bronx, NY, USA. Open Learning, Massachusetts Institute of Technology, Cambridge, MA, USA. Equal contribution (Veronica S. Weiner, David W. Zhou, Pegah Kahali).

## Abstract

During propofol-induced general anesthesia, alpha rhythms undergo a striking shift from posterior to anterior, termed anteriorization. We combined human intracranial recordings with diffusion imaging to show that anteriorization occurs with opposing dynamics in two distinct thalamocortical subnetworks. The cortical and thalamic anatomy involved, as well as their known functional roles, suggest multiple means by which propofol dismantles sensory and cognitive processes to achieve loss of consciousness.

---

Propofol-induced general anesthesia alters arousal, but it is unclear how it disrupts sensory or cognitive processing in humans. Like other GABA-ergic anesthetic drugs, propofol causes widespread slow oscillations (0.1 to 1 Hz), while frontal alpha (8 to 12 Hz) rhythms emerge and the ubiquitous posterior alpha rhythm disappears [1–5]. Slow oscillations are thought to reflect decreased arousal and disrupt cortical function broadly [6]. However, the functional significance of the dual frontal and posterior alpha-band phenomena (termed anteriorization in clinical anesthesiology and neurophysiology [7]) and their circuit architectures within the human brain are not fully understood [8]. Unlike the slow oscillation, alpha dynamics may underlie propofol’s disruptions of sensory and cognitive function, reframing alpha’s ubiquitous role during wakefulness as a sensory processing rhythm. Because the alpha rhythm is generated by thalamocortical mechanisms, anatomical mapping at the cortical and thalamic levels may shed light on the circuitry involved in anteriorization.

We took advantage of two key features of alpha thalamocortical networks to study their brain-wide functional and structural attributes in humans. First, alpha oscillations are coherent within thalamocortical networks, linking thalamic populations with large areas of cortex [9,10]. Therefore, coherence analysis of intracranial recordings offers a way to map the cortical distribution of these oscillatory networks. Second, connected thalamic and cortical regions form system-specific clusters within the thalamocortical network [11], which have been imaged in vivo using probabilistic tractography analysis of diffusion-weighted magnetic resonance images (MRI) [12,13]. Using recordings from hundreds of intracranial channels implanted in eleven epilepsy patients undergoing propofol anesthesia for surgical explantation (Figs. 1a-c), we applied a cross-spectral dimensionality reduction technique [3] to analyze how the spatial structure of coherent alpha rhythms changed during propofol-induced loss of consciousness. Next, we traced fiber connections between each thalamic nucleus and the cortical regions that exhibited changes in alpha-band coherent activity (see *Methods*). This analysis was made possible by collecting a unique dataset from a carefully-orchestrated natural experiment of anesthesia using high-density human intracranial recordings, combined with diffusion images from an open repository.

**Figure 1.**
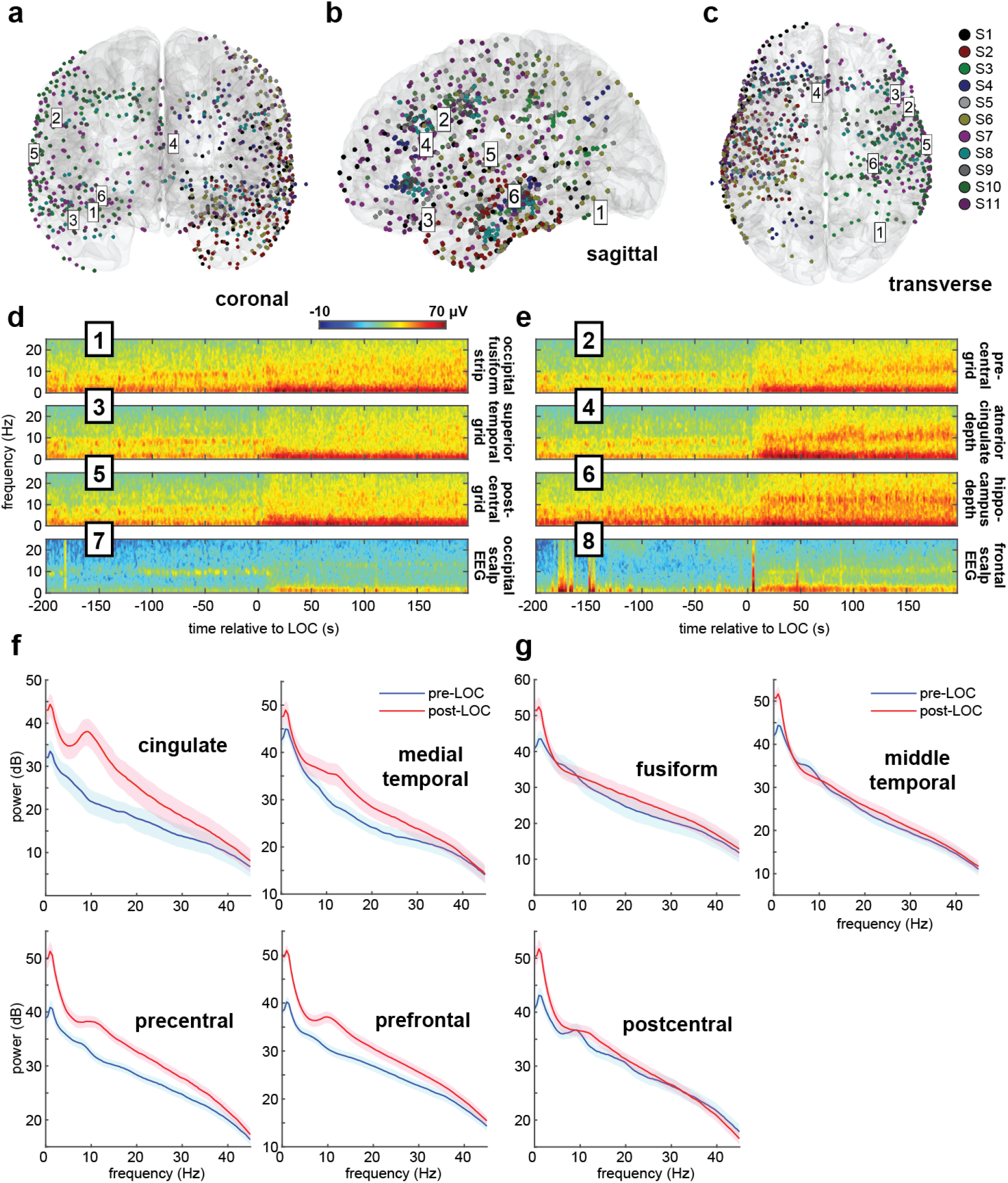
Intracranial electrode coverage and spectral dynamics during propofol-induced general anesthesia. **(a-c)** Intracranial recordings’ (n=897) spatial coverage. Six intracranial channel locations from one subject are labelled. **(d,e)** Multitaper spectrograms in selected channels of a single subject (S7) aligned to LOC time. After LOC, a waking alpha rhythm in posterior regions (odd channels) dissipates, while a broader anesthesia-induced alpha rhythm emerges frontally (even channels). **(f,g)** Pre- and post-LOC multitaper spectra in selected frontal **(f)** and posterior **(g)** regions.

We found that prior to loss of consciousness (LOC), alpha rhythms can be observed in recordings from visual, motor, and auditory cortex (Fig. 1d), consistent with reports of occipital, parietal, and temporal sensory alpha rhythms. After LOC, two rhythms emerge: alpha rhythms appear in cingulate, frontal and medial temporal cortices, and slow wave oscillations appear at all locations (Fig. 1d-e). We determined that anterior and posterior alpha rhythms were spatially-coherent by computing the coherent power spectral density (cPSD) and network-weighted time-frequency signatures from eigendecompositions of the cross-spectral matrix at 10 Hz (see *Methods*). Using cPSD changes between pre-LOC and post-LOC epochs across all recording sites, we identified brain regions in which propofol anesthesia increased or decreased coherent alpha activity (Figs. 2a-c). Propofol achieved its greatest alpha cPSD increases in the cingulate cortex and regions of the frontal lobe, as well as the medial temporal lobe and temporal pole (Fig. 2f). Alpha cPSD was attenuated across the inferior, middle, and superior temporal cortex, as well as in the parietal and occipital cortices. Changes in cPSD magnified regional effects typically seen in shifts of 10-Hz spectral power across the combined channel data, improving the spatial resolution of these cortical maps (Supplementary Figure 3). These coherent alpha dynamics were sharply linked to loss of consciousness in all recordings, as demonstrated by the network-weighted signatures (Figs. 2d-e) of channels coherent at 10 Hz before and after LOC. The post-LOC signature (Fig. 2e) also indicates that propofol’s frontal signature is more broadband than previously thought, spanning both alpha and beta frequencies.

**Figure 2.**
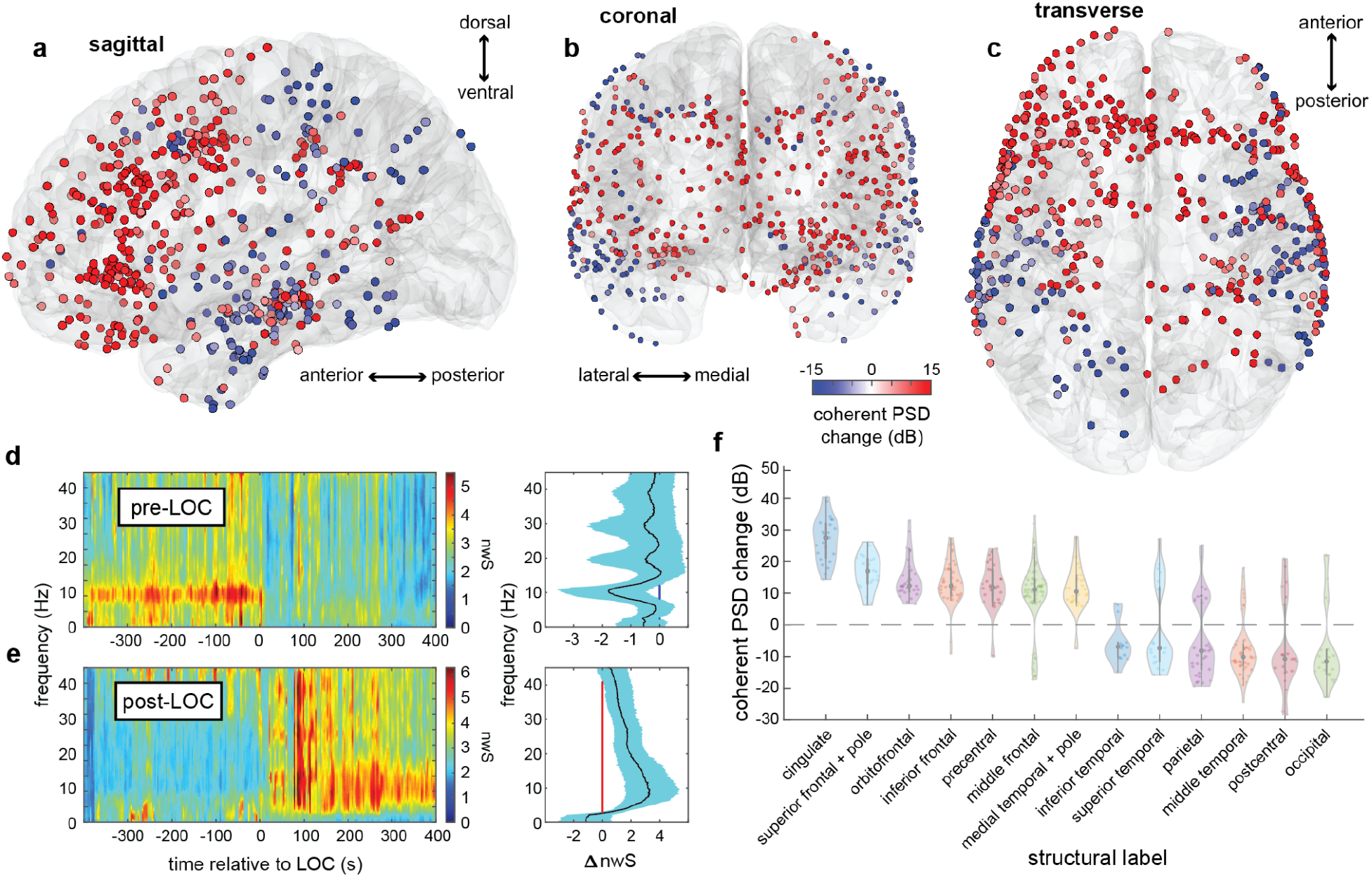
Spatiotemporal mapping of pre- and post-anesthesia coherent alpha networks. **(a-c)** Changes in coherent alpha (10-Hz) cPSD across recording sites in all subjects. **(d,e)** Pre-LOC global coherence principal components depict a narrow 10-Hz rhythm disappearing at LOC (d), and a broader 10-Hz band beginning ~200 seconds after LOC (e). **(f)** cPSD is associated with structural (f) parcellations of brain regions. Frontal midline regions such as cingulate, frontal, and orbitofrontal cortex, as well as medial temporal lobe, show increased alpha-band cPSD after LOC, whereas posterior regions such as somatosensory and visual areas show a decrease. Constituent labels for each structural category are listed in *Supplementary Table 3*.

Confirming previous findings obtained from high-density electroencephalography [4], these results link anterior and posterior coherent alpha networks to known functional divisions of the human cortex situated along the anterior-posterior axis [14,15]. Our results also reveal the cortical distribution of propofol’s anterior alpha signature in greater detail than previous studies by showing that it engages the cingulate and medial temporal cortices. These medial structures, typically inaccessible to measurement by scalp electroencephalography, have not been previously identified as key structural components of propofol’s alpha signature.

Propofol has a number of known biophysical effects that may generate the dual posterior and frontal alpha effects described here. Computational models have suggested multiple mechanisms for thalamic cells [2,8,16] to play a role in propofol’s diverse alpha-band effects. Propofol increases GABA conductance and decay time, which is thought to slow prefrontal thalamocortical circuits to favor alpha oscillations [2,17]. Propofol also inhibits hyperpolarization-activated membrane currents mediated by HCN1 channels, which is believed to disable thalamocortical circuits responsible for posterior alpha waves [8,18,19]. These mechanisms may be linked to functionally specialized thalamic nuclei [20,21].

Several sites in thalamus have been previously studied during propofol anesthesia, but recent evidence for whether propofol increases or decreases alpha synchrony between the cortex and thalamus has been conflicted [5,22–24]. Varying choices of thalamic regions recorded in these studies may explain the inconsistency among their findings regarding propofol’s effects on alpha-band thalamocortical synchrony. To address the controversy, we used probabilistic tractography to determine whether the alpha networks underlying anteriorization might be structurally connected to distinct nuclei within the human thalamus according to their known functional roles (see *Methods*). To do so, we matched our data’s intracranial electrode coordinates with diffusion-weighted MRI data from the WU-Minn Human Connectome Project [25] (see *Methods*).

We found that the posterior alpha network had greater structural connectivity than the anesthesia-induced frontal network to thalamic sensory and sensory association nuclei (Figure 3). Meanwhile, the anesthesia-induced frontal alpha network had greater structural connectivity than the posterior alpha network to thalamic cognitive-, limbic-, and motor-associated nuclei. Some nuclei, such as the centromedian, central lateral and medial pulvinar, appear not to be selectively connected to either network, perhaps explained by their non-specific or diffuse connectivity with cortex (in the case of intralaminar nuclei), or their interconnections with key regions such as the frontoparietal network that span both anterior and posterior cortices. It is known that the sensory alpha is driven by first-order sensory thalamic nuclei such as the lateral geniculate [20] and coordinated by higher-order sensory thalamic nuclei such as the pulvinar [26,27]. Similarly, the prefrontal cortex and the mediodorsal nucleus share connections that support thalamocortical and corticocortical synchrony in the alpha and beta frequency ranges [28–30]. Our results identify distinct groups of nuclei that may synchronize within each oscillatory network, perhaps through corticothalamic feedback [9]. These groups appear to align conspicuously with known systems-specific classes of thalamic nuclei: sensory and sensory association on one hand versus cognitive, limbic, and motor on the other [11].

**Figure 3.**
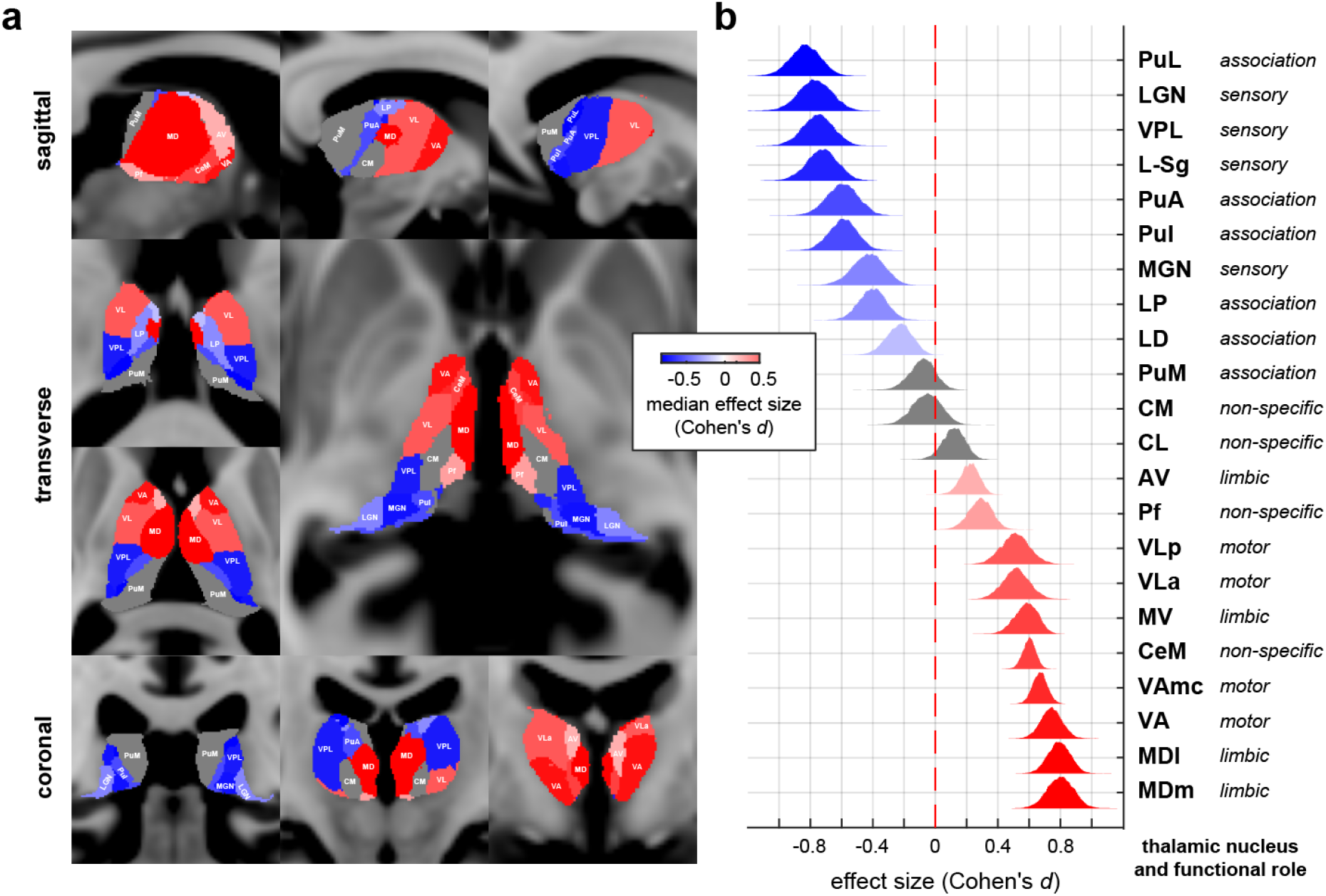
Distinct thalamocortical connections underlying pre- and post-anesthesia alpha networks. **(a)** Thalamic nuclei are selectively connected to distinct networks showing either increased or decreased cPSD after LOC. **(b)** Primary and relay sensory thalamic nuclei selectively connect to alpha cPSD-decreasing regions; nuclei of the executive and cognitive thalamus selectively connect to alpha cPSD-increasing regions. Thalamic nucleus abbreviations: PuL = lateral pulvinar, LGN = lateral geniculate nucleus, VPL = ventral posterolateral, L-Sg = limitans-suprageniculate, PuA = anterior pulvinar, Pul = inferior pulvinar, MGN = medial geniculate nucleus, LP = lateral posterior, LD = laterodorsal, PuM = medial pulvinar, CM = centromedian, CL = central lateral, AV = anteroventral, Pf = parafascicular, VLp = posterior ventral lateral, VLa = anterior ventral lateral, MV = reuniens (medial ventral), CeM = central medial, VAmc = magnocellular ventral anterior, VA = ventral anterior, MDl = parvocellular lateral mediodorsal, MDm = magnocellular medial mediodorsal.

Recent efforts to locate neural correlates of consciousness have made various distinctions between prefrontal and sensory cortex [31–33]. Our results suggest that propofol may impair function in both anterior and posterior networks through distinct mechanisms in order to render sensory information in posterior networks inaccessible to prefrontal areas. Functional mechanisms involving alpha and beta oscillations have been found to gate sensory transmission [34], block perceptual integration of sensory information [35–37], and dysregulate attention [27,38] and working memory [39]. In light of these dynamics, we propose two ways by which propofol may disrupt conscious processing via oscillations: 1) by placing posterior thalamocortical subnetworks in a hyperpolarized state, which degrades feedforward sensory contents and renders them inaccessible to prefrontal oscillatory feedback; and 2) by inducing a non-physiological inhibition of the prefrontal-mediodorsal thalamocortical subnetwork, which restricts its activity to alpha and low-beta bands and interferes with top-down processes. These effects would serve to functionally disrupt feedforward perception on one hand and feedback control of attention, memory and executive function on the other. The two anesthesia-induced alpha network changes that we describe would be predicted to impair both bottom-up and top-down processes, respectively. Disruptions of alpha network dynamics have also been observed in brain states such as traumatic brain injury [40], attention [41], and sleep [42].

Our findings provide empirical evidence that anterior and posterior alpha networks modulated by propofol correspond to known functional subdivisions of the cortex and thalamus. In doing so, our results highlight the need for structural specificity in human studies of anesthesia’s effects on thalamocortical rhythms. Additionally, our results identify locations where propofol may be acting on thalamocortical circuits via GABA or HCN1 mechanisms to either generate or disrupt alpha-band activity [8]. A careful analysis of how propofol compares to other anesthetic drugs with similar or shared mechanisms could further clarify the role that these and other molecular targets play to alter different thalamocortical circuit functions. We note that inhaled ether anesthetics, which also exhibit alpha anteriorization, have GABA and HCN1 mechanisms similar to propofol [43]. In contrast, the anesthetic ketamine is known to act via NMDA and HCN1 mechanisms. Like propofol, ketamine diminishes posterior alpha activity [44], but it neither produces frontal alpha waves nor engages GABA mechanisms in order to disrupt prefrontal functional networks [45,46]. Future animal studies employing multisite recordings of various thalamocortical circuits, such as those connecting the mediodorsal nucleus with prefrontal cortex and the pulvinar nucleus with visual cortex, could shed light on the impact of propofol-induced alpha dynamics on specific sensory and cognitive processes. Overall, our study suggests that propofol acts upon two types of thalamocortical circuits through separate alpha mechanisms to impair functions underlying awareness, distinct from the propofol-induced slow oscillation’s disruption of arousal.

## Online Methods

### Enrollment, demographics, and ethics

We performed intracranial recordings at two hospitals in 14 patients diagnosed with medically-intractable epilepsy following long-term epilepsy seizure monitoring prior to electrode explantation and surgical treatment. Three patients were excluded due to high-amplitude broadband noise across the channel array after LOC. Electrode placement was selected by the patients’ clinicians without regard to study participation. (See Supplementary Table 1 and Figs. 1a-c for channel type and placement.) Patient demographic and clinical information are listed in Supplementary Table 1. All patients gave informed consent in accordance with approved protocols by the hospitals’ Institutional Review Boards. In total, 897 recordings were made from both ECoG arrays (containing strip and grid channels) and depth electrodes, and 99 recordings were made from scalp EEG.

### Anesthesia and behavioral task

Prior to surgery, propofol was administered while subjects performed an auditory behavioral task (Supplementary Figure 1). All subjects received bolus-dose administration. Bolus doses averaged 154.5 mg, with maximum and minimum dosages at 200 and 70 mg, respectively. Two subjects also received infusions after LOC was achieved. Drug doses and protocols were selected by patients’ clinicians without regard to study participation.

Subjects were instructed to perform an auditory button-click task approximately every 4 seconds over a period spanning administration of anesthesia. Responses were presented and recorded using stimulus presentation software (Presentation, Neurobehavioral Systems, Inc., Albany, CA, or EPrime, Psychology Software Tools, Inc., Sharpsburg, PA). The time at which loss of consciousness (LOC) occurred was identified by halving the difference between the last correct behavioral response and the first non-response following a bolus dose of propofol. (See Supplementary Figure 1.) Where accurate responses are unavailable, LOC was assigned to a timepoint 15 seconds following administration of propofol and prior to start of intubation.

### Data acquisition and preprocessing

#### Electrophysiological recordings

Recordings were made during explantation surgery following 1-3 weeks of epilepsy monitoring for detection of epileptogenic foci. Signals were acquired prior to anesthetic induction and continued until after anesthetic induction. EEG and iEEG signals were recorded at 2000, 2500, or 250 Hz depending on acquisition system settings with earlobe (A1/A2), C2, or subdural references when available; otherwise, a common average reference was used. Signals were digitized after applying a high-pass filter above 0.3 Hz (XLTEK, Natus Medical Inc., San Carlos, CA). (See Supplementary Table 1.)

Data were low-pass filtered at 100 Hz using anti-aliasing finite impulse response (FIR) filters, downsampled to 250 Hz, and then notch filtered at 60 Hz and its harmonics and linearly detrended across the entire recording. Data *ν*(*t*) were then re-referenced using an approximate Laplacian (local) reference for intracranial electrodes, using up to *P* = 6 neighbors for each grid electrode and up to *P* = 2 neighbors for strip and depth electrodes using the mean activity in the *P* channels neighboring a channel *x* [Eq1], and then linearly detrended in each 4-second non-overlapping time segment *l* = 1,2,…, *L* [Eq2] where 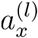 and 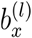 are the segment slope and offset values, respectively:

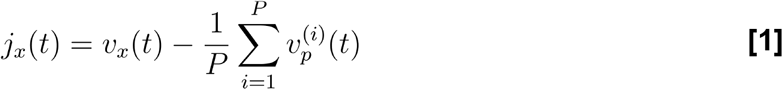

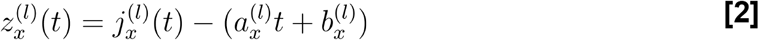

Data segments with missing data or with amplitude greater than 10× the median value across the full recording for each subject were excluded from the global coherence analysis. In individual electrodes, data segments containing epileptiform discharges were excluded from analysis. The total duration of excluded segments totaled less than 5% of recorded data. Three patients were excluded from analysis on the basis of generalized epileptiform discharges in the majority of the channels. No data from the remaining subjects were excluded from the **network-weighted time-frequency representations**.

Two epochs were selected from periods of clean data in each recording and labelled *pre-LOC* and *post-LOC*. Pre-LOC epochs were selected from up to a total of 503 seconds preceding the first propofol dose, and post-LOC epochs were selected from up to a total of 576 seconds following LOC. The beginning of post-LOC epochs were defined by visual inspection of steady-state spectral power in the median spectrogram occuring after paradoxical excitation, denoted by beta (12 to 25 Hz) and gamma band (30 to 60 Hz) power, and prior to burst suppression, which occurred in five subjects. The post-LOC epoch ended at any of these events: a) the first suppression period apparent in the median spectrogram, b) delivery of any anesthetic drug besides propofol, or c) end of the recording. The consistency of spectral patterns in the selected post-LOC epoch was verified by investigators with expertise in EEG data analysis (VSW, DWZ, PLP). (See Supplementary Figure 1 for pre-LOC and post-LOC epoch selections.)

#### Channel coordinate coregistration and morphing

Recording electrodes were identified using preoperative T1-weighted MRI and postoperative CT scans in each patient. RAS coordinates were assigned for all intracranial channels by visual inspection of a maximal intensity projection of the CT, and then projected to the subjects’ individual MRI spaces using coregistration matrices produced using Freesurfer. MRI images were processed by Freesurfer to produce a cortical surface, and channel coordinates were mapped to the nearest surface location using a minimum energy algorithm [47]. Freesurfer was also used to produce structural segmentations and cortical parcellations [48,49], which were visually verified for each electrode and used to label intracranial depth and surface (grid and strip) electrodes, respectively. Poorly situated channels revealed upon verification were removed from the dataset (n = 25, 2.71% of total implanted channels). Coordinates that were not classified by Freesurfer were assigned labels manually against the human atlas by Mai et al [50]. Known anatomical regions were assigned to subsets of the structural labels. Individual MRI images were nonlinearly coregistered to an average brain template in FreeSurfer MNI152 space (cvs_avg35_inMNI152) using Freesurfer’s CVS algorithm [51,52]. Subject-space channel coordinates were morphed to the average brain space using the CVS coregistration outputs and the Freesurfer tool applyMorph, employing a procedure similar to one described in Hamilton et al [53].

#### Patients and channel localization

We performed intracranial recordings from 897 channels in 11 patients as they underwent anesthetic induction with propofol prior to surgical treatment for medically-intractable epilepsy. Patients were asked to perform an auditory behavioral task, and behavioral unresponsiveness was used to define the time of loss of consciousness (LOC). Eight patients ceased task performance within one minute of bolus induction; one patient ceased performance within two minutes, and two patients were excluded from participation in the task by their clinicians. Channels were implanted throughout all lobes of the cortex, including regions of frontal cortex and the visual, auditory, and somatosensory regions of sensory cortex (Fig. 1a-c), as well as white matter (n = 155) and a small number of subcortical regions (brainstem, amygdala, ventral diencephalon, ventricle; n = 8). Locations of electrodes were chosen for epilepsy monitoring and without regard to the current study. Cortical regions sampled by electrode recordings in this study include: caudal and rostral anterior cingulate cortex (n = 17), posterior cingulate cortex (n = 5), caudal middle frontal cortex (n = 26), entorhinal cortex (n = 9), frontal pole (n = 3), fusiform gyrus (n = 33), hippocampus (n = 35), inferior frontal gyrus (n = 53), inferior parietal lobe (n = 14), inferior temporal gyrus (n = 49), medial and lateral orbitofrontal cortex (n = 35), middle temporal gyrus (n = 107), postcentral gyrus (n = 45), precentral gyrus (n = 64), rostral middle frontal cortex (n = 65), superior frontal cortex (n = 19), superior temporal gyrus (n = 63), supramarginal gyrus (n = 44), temporal pole (n = 11).

#### Matched diffusion MRI data

Because high angular resolution diffusion-weighted MRI images were not available for the epilepsy patients enrolled in our study, we assembled a diffusion MRI dataset by matching each non-excluded epilepsy patient to three healthy human surrogates in the 1200-subject WU-Minn Human Connectome Project (HCP) dataset [25]. Surrogates were matched by producing a distance-based ranking of HCP subjects with respect to epilepsy patients using the following demographic and MRI volumetric metrics: age, gender, handedness, brain volume, cortical white matter volume, and thalamic volume. (See Supplementary Table 2.)

The HCP dataset provided high angular resolution diffusion imaging (dMRI) acquired using the 3 Tesla Siemens Skyra “Connectome” scanner. Full dMRI sessions included 6 runs with 3 different gradient tables and oblique axial acquisitions alternating between right-to-left and left-to-right phase encoding directions in consecutive runs. Each gradient table includes approximately 90 diffusion-weighting directions. Diffusion weighting consisted of 3 shells of b=1000, 2000, and 3000 s/mm^2^ interspersed with an approximately equal number of acquisitions on each shell within each run. Six b=0 acquisitions were interspersed throughout each run.

The dMRI data were preprocessed using the HCP diffusion pipeline [54]. The data was further processed with FSL’s BEDPOSTX (Bayesian Estimation of Diffusion Parameters Obtained using Sampling Techniques, modeling crossing X fibers) to model white matter fiber orientations and crossing fibers for probabilistic tractography. BEDPOSTX uses Markov Chain Monte Carlo (MCMC) sampling to build probability distributions on diffusion parameters at each voxel [55].

### Data analysis and statistics

#### Data segmentation

In order to capture the structure of coherent dynamics in the brain states of resting state wakefulness and propofol-induced unconsciousness, we segmented periods of stable oscillatory activity in each patient before and after LOC, labeling these epochs pre-LOC and post-LOC respectively. One of each type of epoch was chosen from each patient’s recording, with pre-LOC epochs beginning up to 513 seconds before LOC and post-LOC epochs terminating up to 775 seconds after LOC. The median length of pre-LOC epochs was 258 seconds (IQR = 280 seconds), and that of post-LOC epochs was 150 seconds (IQR = 216 seconds).

#### Global coherence analysis

To identify frequency-wise coherent networks across the intracranial recording dataset, we used global coherence analysis, a principal component analysis in the frequency domain which identifies the principal modes in the cross-spectral matrix at a frequency *f* [3,56]. For statistical inferences about the output eigenvalues and eigenvectors, we implemented a nonparametric resampling approach for the multitaper cross-spectral matrix using a frequency-domain bootstrap (FDB), which we describe below. We applied eigendecomposition to every bootstrap replicate of the multitaper cross-spectral matrix to obtain empirical distributions of eigenvectors, eigenvalues, coherent power spectral density, and other derivatives of global coherence outputs. Altogether, the eigendecompositions performed at 10 Hz captured on average 41.29% (± 15.58%) of the variance in the pre-LOC data and 48.60% (± 20.64%) in the post-LOC data.

For global coherence analysis, we used *K* = 15 tapers, with non-overlapping time windows of 4 seconds and a half-bandwidth of 2 Hz. In network-weighted time-frequency plots (Eq. 10), we computed spectrograms using the same multitaper parameters with step sizes of 0.1 seconds. In standard multitaper spectrograms, we used K = 3 tapers and 2 second time windows with 0.25 second overlaps. We set *f* = 10 Hz for alpha frequency analyses.

#### Multitaper Cross-Spectrum FDB

We computed tapered Fourier coefficients at frequency *f* for each non-overlapping time series segment *z_x_*(*t*) from a channel *x* ∈ *N* in the channel set *N*, as in:

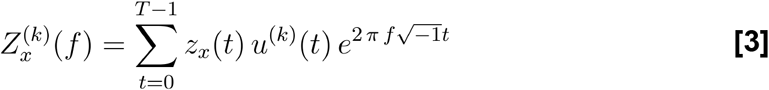

over time *t* = 0,1, … *T* – 1 and *k* = 1,2, … *k*, where *u*^(*k*)^(*t*) is the *k*-th Slepian taper. Then, we computed the |*N*| × |*N*| cross-spectral matrix 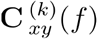 wherein an element in row *x* and column *y* represents the cross-spectrum between channels *x* and *y* for each respective taper and time segment:

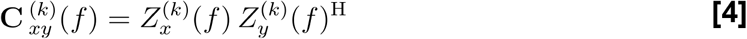

where 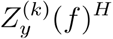 is the complex conjugate transpose of the vectorized tapered Fourier estimates 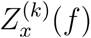.

To generate confidence intervals for cross-spectral estimates, we pursue a frequency-domain bootstrap approach to resampling the epoch-wise multitaper cross-spectral matrix. Proceeding under the assumption that the tapered estimates 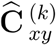 of C_*xy*_ are statistically independent in a given non-overlapping time segment *l* = 1,2,…, *L* of the epoch, we followed the nonparametric approach to resample the average cross-spectrum with replacement [57]. To create a bootstrap replicate 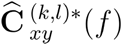 given, *f*, *x*, and *y*, we independently drew K rows and L columns from each matrix of samples 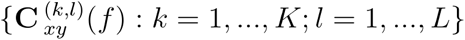. Then, we computed the mean across tapers, and the median across time segments for the real and imaginary components of the multitaper mean separately [58]:

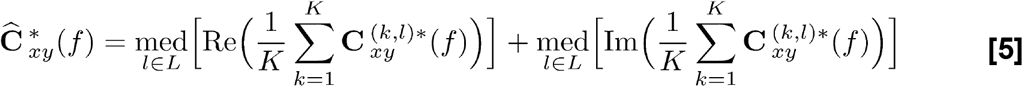

We repeated this procedure to generate *B* = 200 bootstrap replicates of the multitaper cross-spectral matrix per frequency, per epoch (pre-LOC and post-LOC) and per subject. Our resampling procedure can be considered a non-overlapping block bootstrap in the frequency domain.

All subsequent global coherence computations were performed upon each replicate matrix.

#### Cross-spectral matrix eigendecomposition and reconstruction

We performed eigenvalue decompositions (Eq 6) on all replicate cross-spectral matrices [59]:

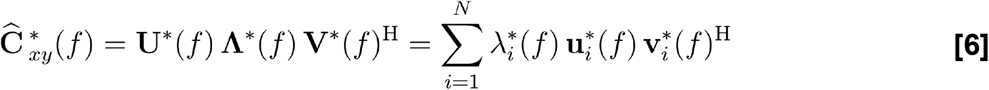

This decomposition represents multi-channel spectral power in terms of orthonormal bases of coherent activity among channels, effectively summarizing the coherent activity at each frequency. In order to estimate the amount of coherent spectral power captured by the principal modes, we reconstructed the cross-spectral matrices using the top three eigenmodes:

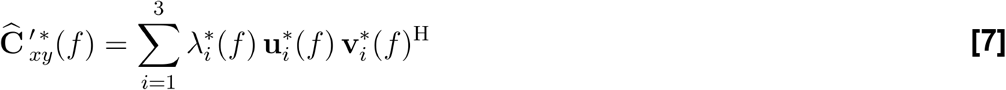

We chose to include the top three eigenmodes to balance parsimony and representation accuracy; we used three eigenmodes across all analyses to ensure consistency. We refer to the diagonal elements 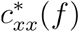 of 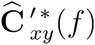 as the ***coherent* power spectral density (cPSD)**.

We estimated the difference in replicated means of the cPSD 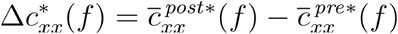 using the bootstrap. We assessed the empirical cumulative distribution functions 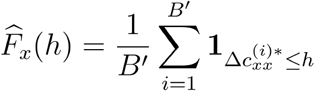 with *B*′ = 10^4^ in order to partition the entire set *N* of intracranial channels into three subsets: 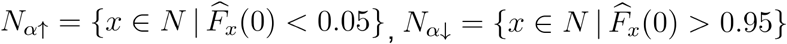, and 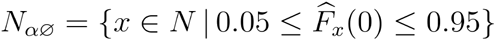, reflecting channels with increasing or decreasing mean coherent power change beyond a 95% confidence level and channels for which the previous two conditions do not apply.

#### Network-weighted time-frequency analysis

To define coherence-based functional networks in a frequency band (such as alpha), we used estimates of 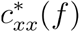 at the respective frequencies *f* = {1, 10, 40} to compute channel weights 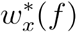 for a given epoch and subject (Eq 8). To visualize the time-frequency activity represented by these network weights over frequencies *f* and time segments *l*, we estimate the **network-weighted spectrum** 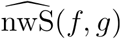, which summarizes the spectrogram across channels ie *i* ∈ *N* using 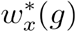 as weights applied to each channel’s spectrogram (Eq 9–10). The number of channels |*N*| is added as a scaling term to allow comparison between data sets with different numbers of channels.

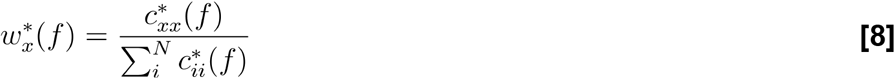

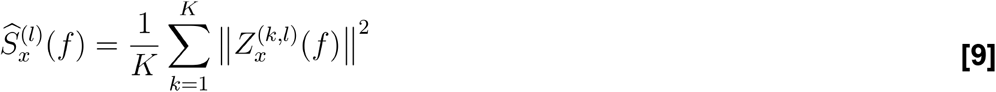

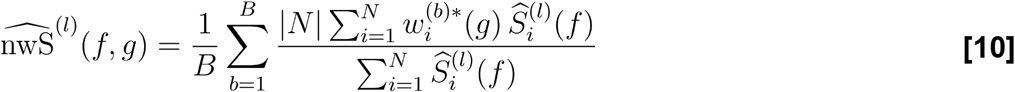

Observe in Eq. 10 that a network represented by a set of equal weights 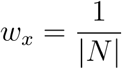 for all channels *x* ∈ *N* would yield a network-weighted time-frequency representation equal to 1 for all *f* and *l*; likewise, time-frequency activity overrepresented by only a few heavily weighted channels in *w_x_*(*g*) would result in values of 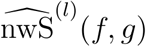 substantially greater than 1. The mean 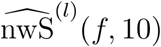 across replicates was computed per subject. In the plots shown in Figure 2, a 2-D median filter with a sliding window of dimensions 4 seconds and 4 Hz was applied to the group-level median network-weighted spectrum. To estimate the overall net change in frequency-domain activity in a network of interest, we computed the group-level mean **network-weighted spectral shift** 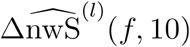 of an alpha network from pre-LOC and post-LOC, using a two-level hierarchical bootstrap of the mean, the first level over subjects (sampled with replacement) and the second level over time segments in each epoch (permuted with replacement per subject-level sample) [60]. Group-level temporal epochs were chosen using all time windows before 0 seconds and after 200 seconds for pre-LOC and post-LOC, respectively, given that each LOC in each subject is aligned at 0 seconds and the majority of time-varying artifacts are attenuated after 200 seconds.

#### Probabilistic tractography

We performed a probabilistic tractography analysis [55] using matched diffusion-weighted MRI data sets from the WU-Minn Human Connectome Project [25]. The intracranial electrophysiology data analysis described above yielded two subsets of channels *N*_*α*↑_ and *N*_*α*↓_ across subjects in which the alpha-band cPSD significantly increased or decreased from pre-LOC to post-LOC, respectively. We used the channel coordinates to produce seed ROIs for tractography analysis. To ensure accurate tracing along white matter tracts, we centered the cortical seed masks at the point on the white matter surface closest to the channel coordinates. For broader sampling of the brain region in the vicinity of each channel, we dilated the seed masks to a 2-mm sphere. To generate thalamic targets, we obtained a probabilistic atlas developed by Iglesias and co-authors [61] and created masks based on 22 nuclei. We used separate masks for the right and left sides of each nucleus, resulting in 44 masks in total. Using FSL’s PROBTRACKX2 (Probabilistic Tracking with Crossing Fibres) [55], we traced streamlines from each channel’s seed mask to all thalamic nuclei using FSL Classification Targets, using 5000 streamlines per voxel in seed masks. We performed tractography analysis for each patient using that patient’s matched samples of diffusion-weighted images.

Each subject from our study was matched with 3 surrogates in the Connectome database according to demographic and brain volumetric indices. ROIs associated with thalamic nucleus membership were defined using a thalamic atlas created by Iglesias and co-authors [61]. We used these thalamic ROIs as targets for traces made from seed coordinates belonging to electrodes in either the frontal or posterior alpha networks.

The specificity of the thalamocortical connection to each nucleus was estimated for the seed masks belonging to the channel sets *N*_*α*↑_ and *N*_*α*↓_ respectively. The number of streamlines reaching each thalamic nucleus from seeds of the channel sets *N*_*α*↑_ and *N*_*α*↓_ was normalized by the total number of streamlines reaching the whole thalamus as well as the size of the given thalamic nucleus (number of voxels). We refer to this quantity as the ROI’s **thalamic nucleus connectivity**:

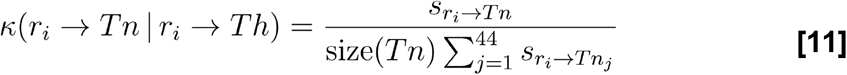

where *s*_*r*→*Tn*_ is the number of total streamlines between a seed mask ROI *r* belonging to channel *i* ∈ *N*_*α*↑_ or *N*_*α*↓_ and the mask of a thalamic nucleus *Tn*, and where *Th* is the union of all thalamic nucleus masks.

We wondered whether streamlines from a cortical ROI to a contralateral thalamic nucleus would be anatomically plausible. In primary-level sensory thalamocortical pathways, they are considered not to be; however, there is evidence in non-human primates that frontal areas may connect bilaterally [62,63]. Therefore, we calculated two versions of *κ*: one in which each *Tn* is the union of left and right nucleus masks, and another in which each *Tn* is defined strictly as the nucleus ipsilateral to the respective cortical ROI *r_i_*.

To infer the connection preference of a given thalamic nucleus to regions covered by either *N*_*α*↑_ or *N*_*α*↓_, we calculated the empirical probability distributions of the effect size (Cohen’s *d*) between thalamus nucleus connectivity values of ROIs in and *N*_*α*↑_ and *N*_*α*↓_:

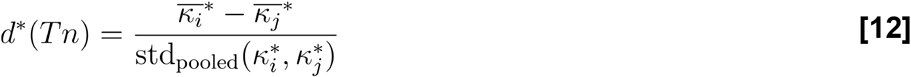

where
*κ_i_* ∈ {*κ*(*r_i_* → *Tn*| *r_i_* ∈ *Th*) ∀*i* ∈ *N*_*α*↑_} and *κ_j_* ∈ {*κ*(*r_j_* → *Tn*| *r_j_* ∈ *Th*) ∀*j* ∈ *N*_*α*↓_}. We resample with the same two-level hierarchical technique described above. Figure 4B uses the bilateral version of *κ*. (See Supplementary Figure 4 for the ipsilateral version.)

## Acknowledgements

This work was supported by NIH grants P01GM118269 (Brown), 1R01AG056015 (Purdon), R21DA048323 (Purdon), T32EB019940 (Zhou), and the Tiny Blue Dot Foundation (Purdon). Data were provided in part by the Human Connectome Project, WU-Minn Consortium (1U54MH091657 (Van Essen, Ugurbil)) funded by the 16 NIH Institutes and Centers that support the NIH Blueprint for Neuroscience Research; and by the McDonnell Center for Systems Neuroscience at Washington University. We would like to thank Bram Diamond for the 1mm isotropic MNI atlas of thalamic nuclei. We are grateful to Sourish Chakravarty, Brian Edlow, Michael Halassa, Nancy Kopell, Michelle McCarthy, Samuel Snider, and Carmen Varela for their valuable comments and suggestions. Above all, we are deeply indebted to the volunteers who contributed their time and efforts as surgical patients to our research.

## Author contributions

List of author initials: VSW, DWZ, PK, EPS, RAP, LA, MDS, ENE, AFS, ALS, SSC, ENB, PLP VSW, ENE, SSC, ENB, PLP conceived and planned the experiments.

VSW, RAP, LA, MDS, ENE, AFS, ALS, SSC, PLP carried out the experiments.

VSW, DWZ, PK, PLP preprocessed the data.

VSW, DWZ, PK wrote the preprocessing and analysis software.

VSW, DWZ, PK, EPS, PLP performed the data, statistical, and computational analyses and wrote mathematical descriptions.

VSW, DWZ, PK, PLP drafted the manuscript.

VSW, DWZ, PK, EPS, PLP reviewed and edited the manuscript.

VSW, DWZ, ENB, PLP funded the project.

## Data materials and availability

Diffusion-weighted imaging data is publicly available at https://www.humanconnectome.org/study/hcp-young-adult. Code will be made available in a public repository prior to publication.

## Supplementary Tables and Figures

**Supplementary Table 1.**
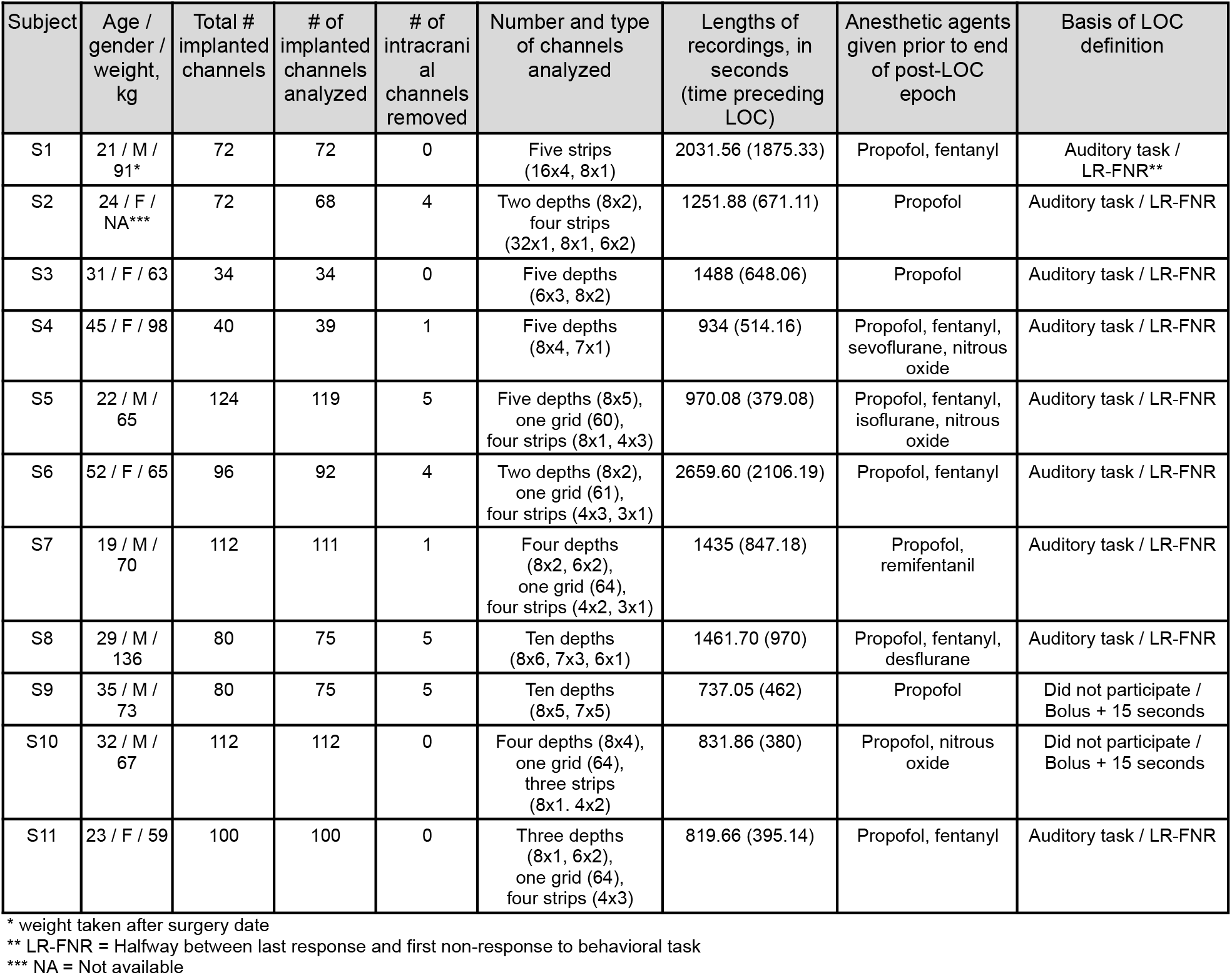
Patient demographic and clinical information.

| Subject | Age / gender / weight, kg | Total # implanted channels | # of implanted channels analyzed | # of intracranial channels removed | Number and type of channels analyzed | Lengths of recordings, in seconds (time preceding LOC) | Anesthetic agents given prior to end of post-LOC epoch | Basis of LOC definition |
| --- | --- | --- | --- | --- | --- | --- | --- | --- |
| S1 | 21 / M / 91* | 72 | 72 | 0 | Five strips (16x4, 8x1) | 2031.56 (1875.33) | Propofol, fentanyl | Auditory task / LR-FNR** |
| S2 | 24 / F / NA*** | 72 | 68 | 4 | Two depths (8x2), four strips (32x1, 8x1, 6x2) | 1251.88 (671.11) | Propofol | Auditory task / LR-FNR |
| S3 | 31 / F / 63 | 34 | 34 | 0 | Five depths (6x3, 8x2) | 1488 (648.06) | Propofol | Auditory task / LR-FNR |
| S4 | 45 / F / 98 | 40 | 39 | 1 | Five depths (8x4, 7x1) | 934 (514.16) | Propofol, fentanyl, sevoflurane, nitrous oxide | Auditory task / LR-FNR |
| S5 | 22 / M / 65 | 124 | 119 | 5 | Five depths (8x5), one grid (60), four strips (8x1, 4x3) | 970.08 (379.08) | Propofol, fentanyl, isoflurane, nitrous oxide | Auditory task / LR-FNR |
| S6 | 52 / F / 65 | 96 | 92 | 4 | Two depths (8x2), one grid (61), four strips (4x3, 3x1) | 2659.60 (2106.19) | Propofol, fentanyl | Auditory task / LR-FNR |
| S7 | 19 / M / 70 | 112 | 111 | 1 | Four depths (8x2, 6x2), one grid (64), four strips (4x2, 3x1) | 1435 (847.18) | Propofol, remifentanyl | Auditory task / LR-FNR |
| S8 | 29 / M / 136 | 80 | 75 | 5 | Ten depths (8x6, 7x3, 6x1) | 1461.70 (970) | Propofol, fentanyl, desflurane | Auditory task / LR-FNR |
| S9 | 35 / M / 73 | 80 | 75 | 5 | Ten depths (8x5, 7x5) | 737.05 (462) | Propofol | Did not participate / Bolus + 15 seconds |
| S10 | 32 / M / 67 | 112 | 112 | 0 | Four depths (8x4), one grid (64), three strips (8x1, 4x2) | 831.86 (380) | Propofol, nitrous oxide | Did not participate / Bolus + 15 seconds |
| S11 | 23 / F / 59 | 100 | 100 | 0 | Three depths (8x1, 6x2), one grid (64), four strips (4x3) | 819.66 (395.14) | Propofol, fentanyl | Auditory task / LR-FNR |

\* weight taken after surgery date
\*\* LR-FNR = Halfway between last response and first non-response to behavioral task
\*\*\* NA = Not available

**Supplementary Table 2.** Patient-specific analysis metadata.

| Subject | Length of pre-LOC coherence analysis epochs in seconds | Length of post-LOC coherence analysis epochs in seconds | Percent variance explained in top three pre-LOC principal component (bootstrap median, IQR) | Percent variance explained in top three post-LOC principal component (bootstrap median, IQR) |
| --- | --- | --- | --- | --- |
| S1 | 345.77 | 71.97 | 38.87, 2.43 | 55.26, 21.41 |
| S2 | 150.00 | 419.00 | 42.47, 2.33 | 40.12, 1.22 |
| S3 | 461.46 | 118.90 | 71.56, 1.37 | 68.74, 2.31 |
| S4 | 503.17 | 100.00 | 63.75, 1.36 | 78.23, 5.47 |
| S5 | 258.08 | 190.80 | 38.26, 2.753 | 33.00, 2.60 |
| S6 | 150.00 | 103.80 | 23.26, 1.50 | 20.9, 1.18 |
| S7 | 100.00 | 475.91 | 28.56, 1.53 | 23.10, 0.98 |
| S8 | 410.80 | 324.30 | 38.97, 1.79 | 54.74, 1.87 |
| S9 | 377.90 | 197.80 | 46.39, 1.40 | 76.41, 2.50 |
| S10 | 44.20 | 150.00 | 42.41, 14.08 | 54.35, 6.20 |
| S11 | 195.14 | 101.00 | 19.57, 0.76 | 29.79, 1.59 |

**Supplementary Table 3.** Categorization of structural regions and labels.

| Structural region category | Freesurfer structural labels |
| --- | --- |
| cingulate | rostral anterior, caudal anterior, posterior gyri, isthmus of |
| superior frontal + pole | frontal pole and superior frontal gyrus |
| orbitofrontal | medial and lateral orbitofrontal cortex |
| inferior frontal | pars orbitalis, pars opercularis, pars triangularis |
| precentral | precentral gyrus |
| middle frontal | middle frontal gyrus |
| medial temporal + pole | temporal pole, hippocampus, parahippocampal, entorhinal cortices |
| inferior temporal | inferior temporal gyrus |
| superior temporal | superior temporal gyrus |
| parietal | superior and inferior parietal lobules |
| middle temporal | middle temporal gyrus |
| postcentral | postcentral gyrus |
| occipital | superior occipital cortex, precuneus, pericalcarine, fusiform, lingual gyri |

**Supplementary Table 4.** Human Connectome Project matches.

| Subject | Human Connectome Project matched subject IDs |
| --- | --- |
| S1 | 107422, 187850, 573451 |
| S2 | 140925, 480141, 565452 |
| S3 | 136732, 175237, 406836 |
| S4 | 202820, 757764, 905147 |
| S5 | 107422, 187850, 573451 |
| S6 | 202820, 757764, 905147 |
| S7 | 107422, 187850, 573451 |
| S8 | 346137, 594156, 958976 |
| S9 | 108222, 540436, 592455 |
| S10 | 173738, 397861, 818455 |
| S11 | 205725, 706040, 877269 |

**Supplementary Figure 1.**
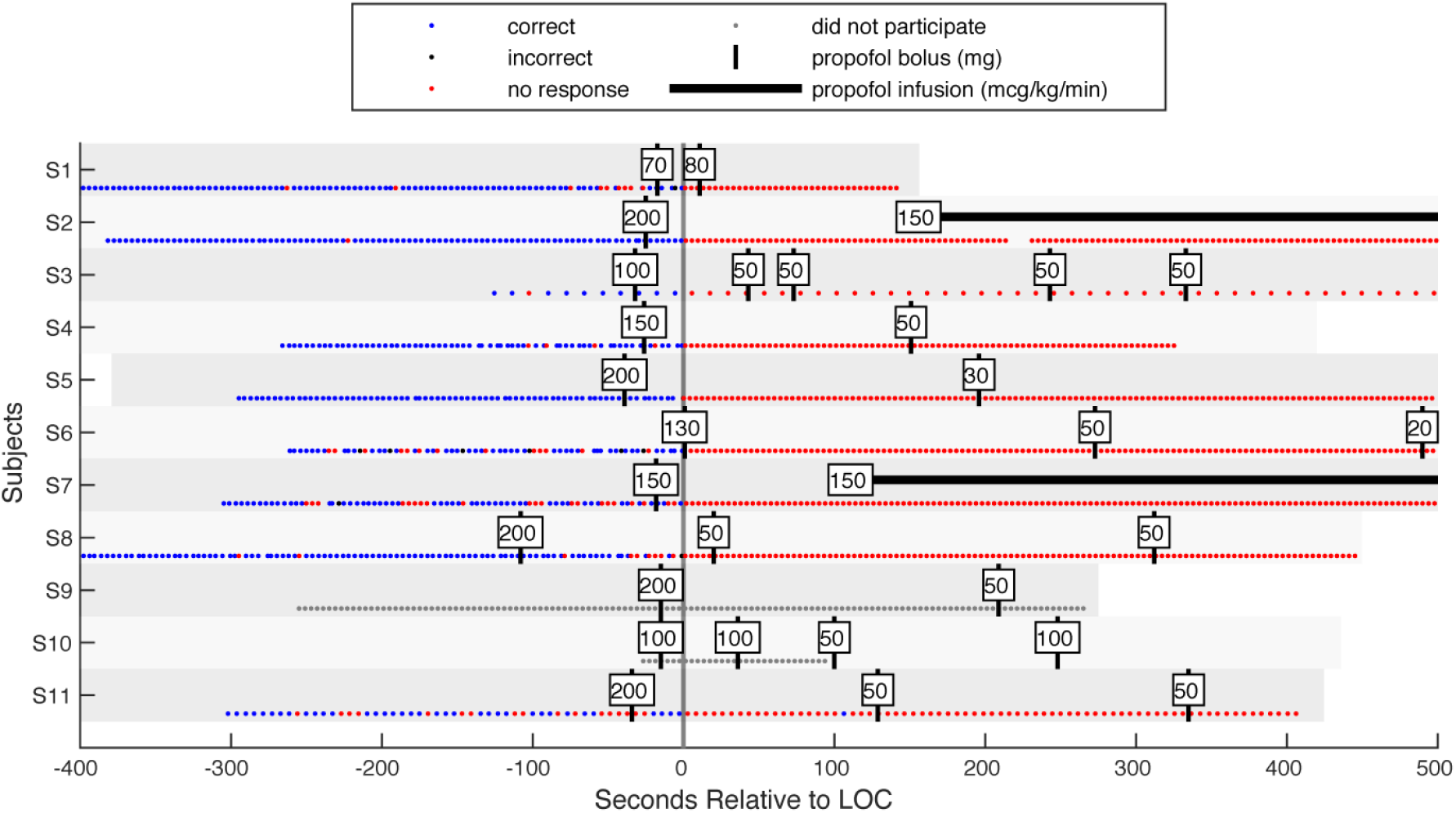
Patient anesthesia administration time courses. Shaded gray rows cover the start to end of each recording. Two patients did not participate in the behavioral paradigm, due to either failure to execute the task or exclusion at the request of the clinician. For these two patients, LOC was marked at 15 seconds after first propofol bolus injection.

**Supplementary Figure 2.**
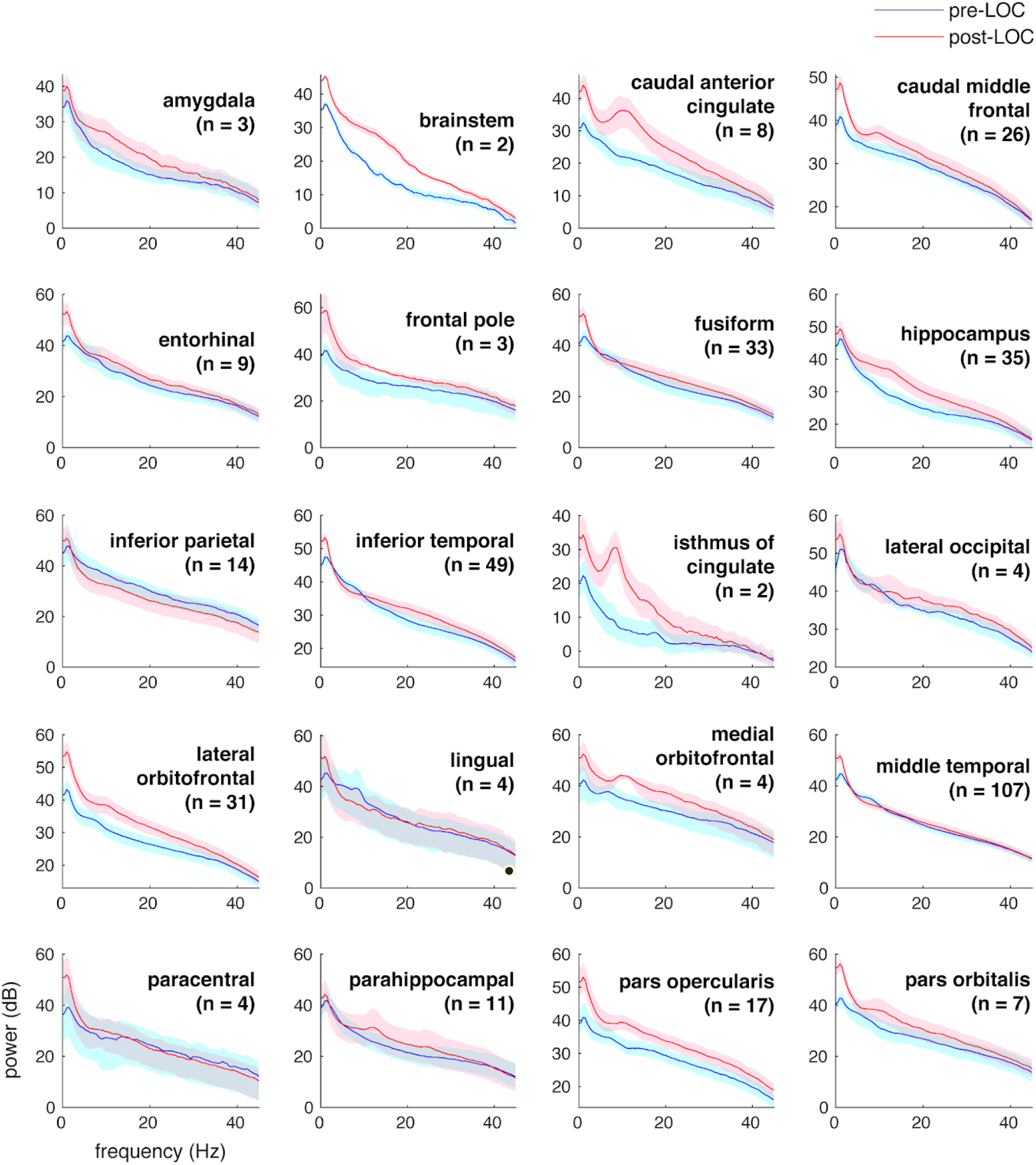
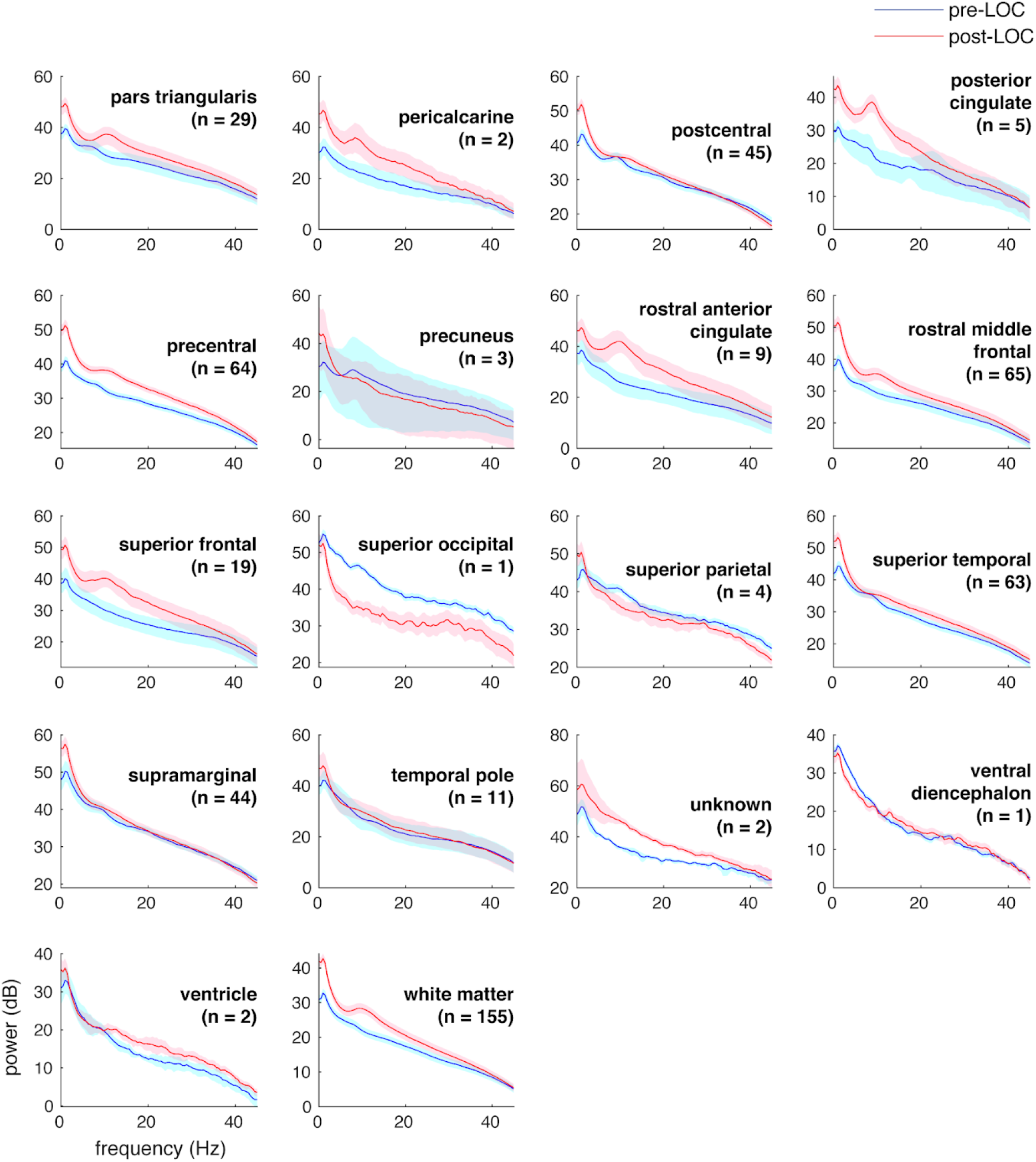
Pre- and post-LOC spectra across recorded regions. Pre- and post-LOC multitaper spectra from all anatomical regions that contained implanted channels. Shaded boundaries represent the 95% bootstrap standard error of the mean.

**Supplementary Figure 3.**
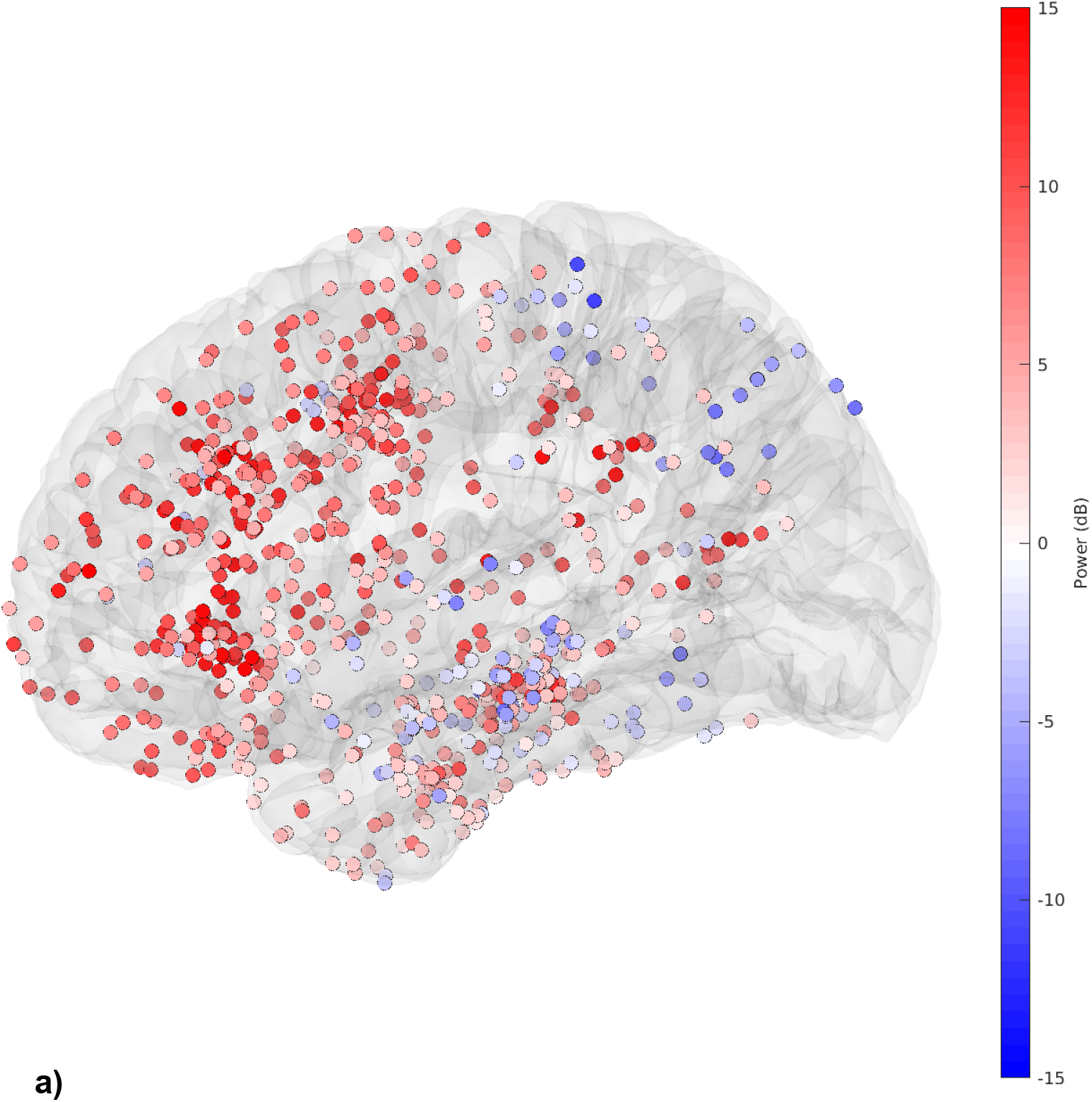
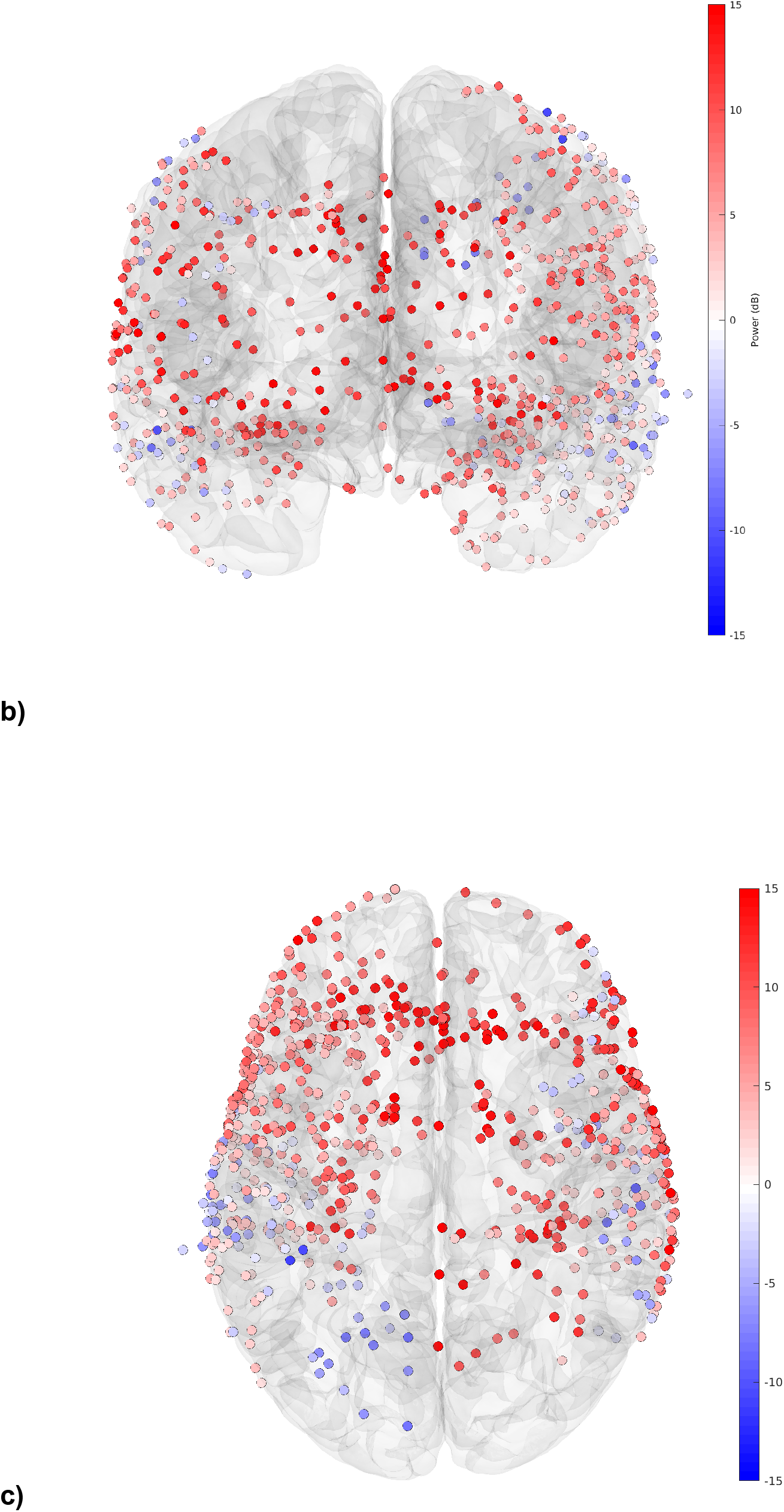
Alpha power changes across channels. Channel-wise alpha (10 Hz) multitaper spectral power changes between pre-LOC and post-LOC epochs, in sagittal (**a**), coronal (**b**), and transverse (**c**) views. Color intensities represent median bootstrap differences (similar to Figs. 2a-c) between 10-Hz multitaper spectral power in each epoch.

**Supplementary Figure 4.**
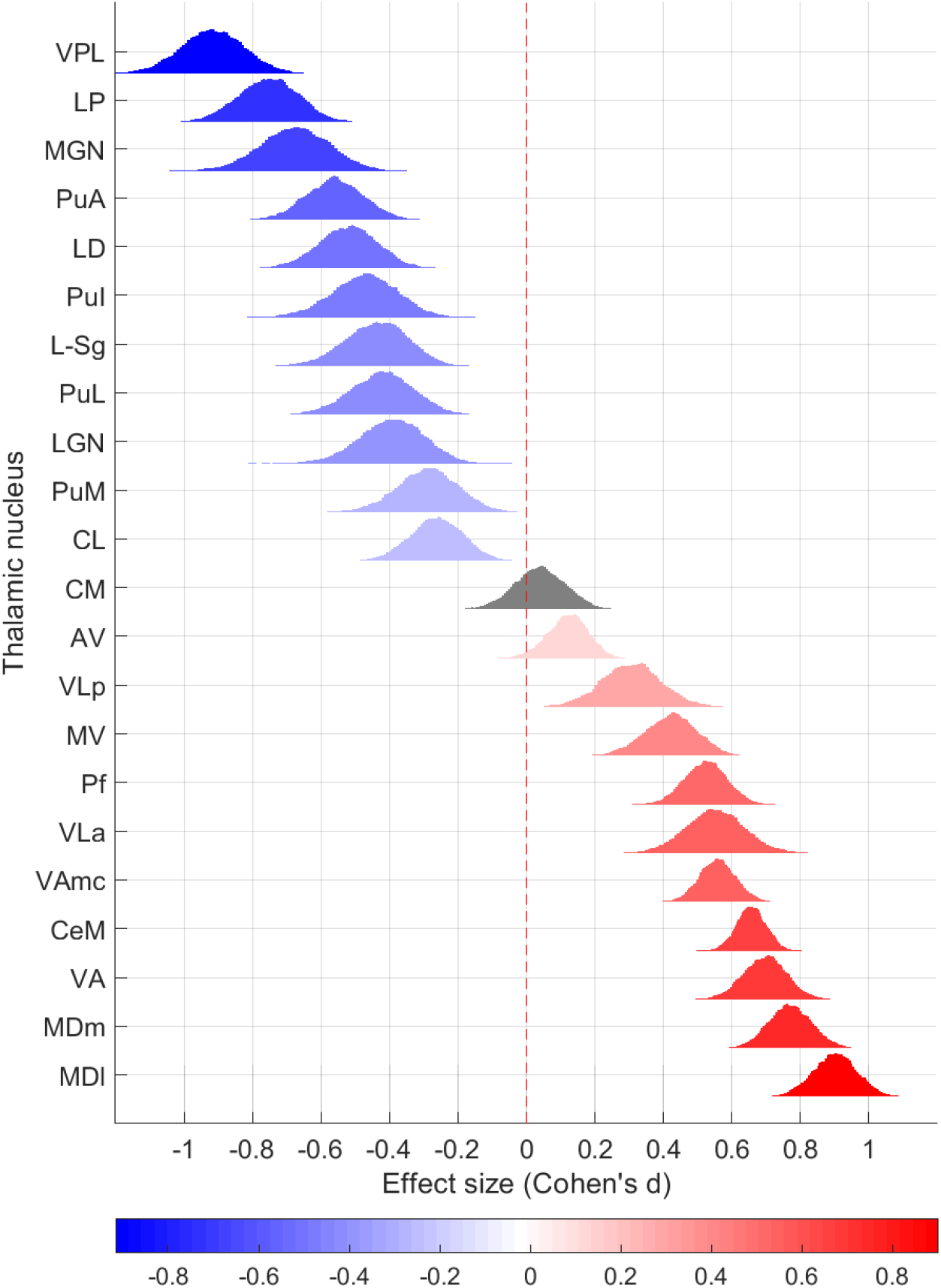
Selective connectivity of ipsilateral thalamocortical tracts. Re-analysis of preferential connectivity to alpha-increasing and -decreasing regions (Fig. 4b), using ipsilateral thalamus-to-cortex streamlines only. (See Methods.)

